# Benchmark Problems for Dynamic Modeling of Intracellular Processes

**DOI:** 10.1101/404590

**Authors:** Helge Hass, Carolin Loos, Elba Raimundez Alvarez, Jens Timmer, Jan Hasenauer, Clemens Kreutz

**Author notes:** These authors contributed equally (Helge Hass, Carolin Loos).

## Abstract

**Motivation:** Dynamic models are used in systems biology to study and understand cellular processes like gene regulation or signal transduction. Frequently, ordinary differential equation (ODE) models are used to model the time and dose dependency of the abundances of molecular compounds as well as interactions and translocations. A multitude of computational approaches have been developed within recent years. However, many of these approaches lack proper testing in application settings because a comprehensive set of benchmark problems is yet missing.

**Results:** We present a collection of 20 ODE models developed given experimental data as benchmark problems in order to evaluate new and existing methodologies, e.g. for parameter estimation or uncertainty analysis. In addition to the equations of the dynamical system, the benchmark collection provides experimental measurements as well as observation functions and assumptions about measurement noise distributions and parameters. The presented benchmark models comprise problems of different size, complexity and numerical demands. Important characteristics of the models and methodological requirements are summarized, estimated parameters are provided, and some example studies were performed for illustrating the capabilities of the presented benchmark collection.

**Availability:** The models are provided in several standardized formats, including an easy-to-use human readable form and machine-readable SBML files. The data is provided as Excel sheets. All files are available at https://github.com/Benchmarking-Initiative/Benchmark-Models, with MATLAB code to process and simulate the models.

## 1 Introduction

Dynamic models based on ordinary differential equations (ODEs) have become a widely used tool in systems biology to quantitatively describe regulatory processes in living cells. Within this approach, known biochemical interactions of important compounds can be translated into rate equations describing the temporal evolution of the state of biological processes. Experimental data is then used to estimate parameters like rate constants or initial concentrations and to validate or improve the model structure.

The dimensionality and nonlinearity of these models constitute a challenge for numerical and statistical methods regarding parameter estimation and identification of the most plausible model structure. For that reason, a multitude of new modeling techniques have been developed within recent years. However, they are often not well-tested especially in realistic application settings and therefore performance benefits or limitations are unknown (Vyshemirsky and Girolami, 2008; Lillacci and Khammash, 2010; Raue *et al*., 2013; Hug *et al*., 2013; Degasperi *et al*., 2017; Maier *et al*., 2016; Stapor *et al*., 2018). Since the performance of computational approaches depends on model characteristics such as nonlinearity, number of parameters or amount of experimental data, it is essential to have a reasonably large set of benchmark problems. These need to cover a broad range of application settings in order to generalize results obtained in performance studies to new modeling projects.

One frequent limitation is that realistic measurements are typically not available for evaluations. Simulated data, as an example, is often much more informative in terms of number of data points (Tönsing *et al*., 2014) and does not have a complex noise structure (Villaverde *et al*., 2015) like measurements from living cells. Moreover, in most cases experimental measurements require augmenting the equations of the dynamic model with so-called observation functions containing scalingsand/or offset parameters, together with transformations of the data such as a log-transformation.

In many scientific fields benchmark collections are available, however, only a limited number of benchmark problems are currently available for modeling intracellular processes and they cover only a small set of application setups: (i) Six benchmark models have been published by Villaverde *et al*. (2015), however for most of them, only simulated data are provided. For the models with experimental data, one has less data points than parameters, and the other provides its equations only in a compiled version, which limits their use for model evaluation. (ii) Additional benchmark problems were defined within the DREAM6 (Dialogue on Reverse-Engineering Assessment and Methods) and DREAM7 challenges. However, both challenges only had simulated data available because the models do not represent real biological networks occurring in specific living cells. In addition, abundances of the molecular compounds were assumed as known initial values and the dynamic variables were assumed as directly measured without observation functions which renders these problems as rather unrealistic. (iii) Public repositories, e.g., the *Biomodels database* (Le *Novere et al*., 2006) provide a large number of realistic / published models. Unfortunately, for most models the measured data used for calibration is not or only partly provided. Moreover, if the data is published, the description of the link between model and data is often not sufficient, i.e., the noise model and observation functions are not comprehensively defined as required for a non-ambiguous benchmark problem. One major reason for this might be that current standards for defining models like the Systems Biology Markup Language (SBML) (Hucka *et al*., 2003) only comprise the biological part of the model but do not contain equations for observations and noise models used to estimate parameters. Standards for the encoding of experimental descriptions, such as the Simulation Experiment Description Markup Language (SED-ML) (Waltemath *et al*., 2011), are unfortunately not yet used widely and only supported by a fraction of the available tools.

In this manuscript, 20 models of biochemical reaction networks are presented which should serve as a comprehensive set of benchmark problems enabling testing of a multitude of data-based modeling approaches. The models have different complexity ranging from 9 to 269 parameters. All models comprise measured data (21 to 27132 data points per model). We also provide measurement errors either determined experimentally or from an underlying error model.

## 2 Methodology

### 2.1 Pathway models

Biochemical reaction networks can be modeled using reaction rate equations,

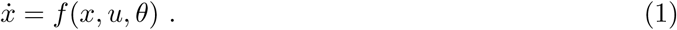

which describe the dynamics of compound concentrations 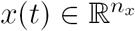 as a function of parameters *θ* (Section 2.3) and inputs 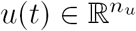 (Section 2.4).

The initial values *x*(0) of Eq. (1) might be known. However, in most applications some elements of *x*(0) are unknown and defined as parameters, i.e., 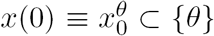, or functions of parameters, i.e., *x*(0) *≡ x*_0_(*θ*). Mathematically, we distinguish between three classes:

1. The initial conditions might be known / given, e.g., zero before treatment.
2. The initial conditions might be analytical functions of the parameters, e.g., analytical solutions to a steady-state constraint (Rosenblatt *et al*., 2016).
3. The initial conditions might be non-analytical expressions of the parameters, e.g., the result of a pre-simulation 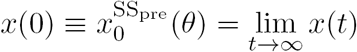 of an experimental condition (Rosenblatt *et al*., 2016; Fiedler *et al*., 2016).

For a detailed discussion we refer to Rosenblatt *et al*. (2016) and Fiedler *et al*. (2016).

### 2.2 Measurement errors

The state variables of reaction rate equations are linked to measurements via observation functions *g*_*i*_(*x, θ*), *i* = 1*,…, N*_obs_, which describe the properties of the experimental device / technique used to acquire measurement data. The observation functions might be nonlinear functions of the state variables, e.g., if the readout saturates, for considering detection limits, and comprise scalings (Loos *et al*., 2018). For all presented benchmark models, independent normally distributed, additive errors are assumed for the measurements

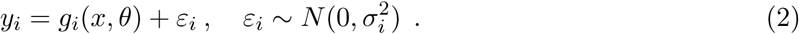

Note that in the chosen notation, index *i* enumerates each observation/data point *y*_*i*_ at a specific time point and each corresponding standard deviation *σ*_*i*_ of the measurement error individually.

We consider two broad classes of error models:

1. The standard deviation *σ*_*i*_ of measurement errors might be determined as part of the experiment and processing of raw data, e.g., by computing standard errors across replicates. In this case, each data point *y*_*i*_ has a given, fixed value *σ*_*i*_ specifying the accuracy of the measurement.
2. Standard deviations might be unknown and therefore described as error models with error parameters which might be jointly estimated with other model parameters. The function can depend on parameters, state variables or both.

While class 1 yields a parameters estimation problem with fewer parameters, class 2 does not require the calculation of *σ*_*i*_ from a potentially small number of replicates and the statistical model accounts for imperfect knowledge of *σ*_*i*_ (Raue *et al*., 2013).

An error model *E* describes the dependence of the standard deviation of an observation on the error parameters *θ*_err_ and the state variables *x*, *σ*_*i*_ = fnc(*g*_*i*_(*x, θ*)*, θ*_err_). The most basic parameter-dependent error models are unknown standard deviations for the individual observations, *∀i*: *σ*_*i*_ *≡ θ*_abserr*,i*_, or sets of observations *I*_*s*_, *s* = 1*,…, n*_*s*_, i.e.,

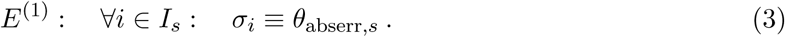

Parameter- and state-dependent error models are for instance

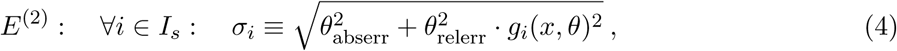

and

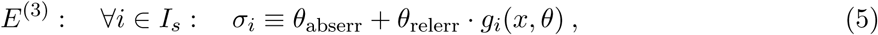

with two parameters for absolute or relative noise levels. *E*^(2)^ is obtained if relative and absolute errors are assumed as two independent sources of variability. *E*^(3)^ is a phenomenological model which often realistically describes absolute and relative components of observed measurement errors.

### 2.3 Parameters

Dynamic models in systems biology comprise up to three classes of parameters:

- Dynamic parameters *θ*_dyn_ that determine the initial states *x*(0) and the dynamics of the process, see Eq. (1). These parameters are rate constants such as association/dissociation rates or -constants, translocation rates between intra- or extracellular compartments, or parameters like Michaelis-Menten- and Hill-coefficients, efficiencies of genetic perturbations or parameters of input functions. We note that the dynamic parameters *θ*_dyn_ do not change over time, although the name might suggest otherwise.
- Observation parameters *θ*_obs_ that describe the relationship between concentrations of intracellular compounds with outputs, e.g., intensities in an assay. These parameters are for example scaling factors or offsets (Weber *et al*., 2011).
- Error parameters *θ*_err_ that describe the unknown noise levels (see Section 2.2).

Since dynamic parameters depend on the biological context and observationand error parameters are determined by the experimental setup, there is often only a limited amount of prior knowledge about parameters available. For the benchmark models, upperand lower bounds are defined for all parameters. In most cases, these bounds cover eight orders of magnitude or even more. In some cases, additional prior knowledge in terms of prior distributions or penalties is available for specific parameters.

The parameters of the biological process are often transformed to improve the convergence of optimization (Raue *et al*., 2013) and to eliminate structural non-identifiabilities (Maiwald *et al*., 2016). A common practice is the transformation of the parameters from linear to logarithmic scale. However, there are also problem-specific transformations as described in the Supplement for the Bachmann or Becker models.

### 2.4 Inputs

Inputs *u* describe the dependence of the biochemical reaction network on external factors as well as perturbations. Examples are externally controlled concentration of ligands or nutrients, or genetic perturbations like knockouts or overexpression. Time dependent inputs are often parameterized functions such as polynomials, splines (Schelker *et al*., 2012) or control vectors (Banga *et al*., 2005). Time-dependent inputs *u ≡ u*(*t*) might depend on parameters which is denoted by *u*(*t, θ*) in the following.

## 3 Model and data formats

For a thorough evaluation of computational methods, we provide a set of 20 published models and corresponding data sets. The models have been extracted from the literature and have been developed by more than 10 different research groups. The information is stored in an easily accessible and standardized format, including an Excel file with general specifications of the model and its fit results. Measurements and model equations are stored as separate Excel files and for each experiment individually. In the data files, measurements with uncertainties and results from the corresponding model simulations are stored. The model files contain finalized ODEs including experiment-specific parameter assignments and observation functions, and are provided as user-readable Excel file and in the standardized, machine-readable SBML standard (Hucka *et al*., 2003). For a detailed description of the provided files, we refer to Supplementary Section 1.

## 4 Results

### 4.1 Benchmark collection

The main focus of this paper is to introduce a comprehensive collection of benchmark problems and their formulation in a standardized format. A comprehensive overview of the benchmark problems is provided in Table 1 on page 12.

**Table 1:** Table summarizing the 20 benchmark models and their properties. The models are abbreviated with the last name of the first author. Many models are based on Western blot data. The number of experimental conditions is specified as the number of different simulation conditions. The feature abbreviations denote the following: C = several compartments, *E*^(1)^ = constant error parameters, Eq. (3), *E*^(2)^ = error model of Eq. (4), *E*^(3)^ = error model of Eq. (5) of main manuscript, Ex = known measurement errors, ev = events, NI = non-identifiable parameters, *u*(*t*) = time dependent input function, *u*(*t, θ*) = input function with unknown parameter(s). Initial values are specified according to the following order: 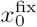 = known initial values 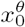 = initial condition given by unknown parameters, *x* (*θ*) = parameter dependent functions, and 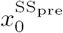 = pre-equilibration for initial steady state conditions. The models are described in more detail in Supplementary Section 3.

| Name | Description | Biochemical species | Observables | Data points | Experimental conditions | Parameters | Features |
| --- | --- | --- | --- | --- | --- | --- | --- |
| Bachmann | The model by Bachmann *et al.* (2011) describes JAK2/STAT5 regulation via two transcriptional negative feedbacks, CIS and SOCS3 in murin blood forming cells | 25 | 6 | 542 | 23 | 113 | C, $E^{(1)}$, NI, $x_0^\theta$ |
| Becker | The model by Becker *et al.* (2010) shows that rapid EpoR turnover and large intracellular receptor pools enables linear ligand response. | 6 | 4 | 85 | 13 | 16 | $E^{(1)}$, $x_0(\theta)$ |
| Beer | The model by Beer *et al.* (2014) uses *Escherichia coli* as chassis to demonstrate heterologous T domain exchange in non-ribosomal peptide synthetases (NRPSs). | 4 | 2 | 27132 | 19 | 72 | $E^{(1)}$, ev, $u(t, \theta)$, $x_0^\theta$ |
| Boehm | The model by Boehm *et al.* (2014) evaluates possible homo- and heterodimerization of the transcription factor isoforms STAT5A and STAT5B. | 8 | 3 | 48 | 1 | 9 | C, $E^{(1)}$, $u(t, \theta)$, $x_0(\theta)$ |
| Brännmark | The model by Brännmark *et al.* (2010) describes insulin signaling in adipocytes. | 9 | 2 | 43 | 8 | 22 | $E^{(1)}$, ev, NI, $u(t)$, $x_0^{\text{SSpre}}(\theta)$ |
| Bruno | The model by Bruno *et al.* (2016) investigates the activity of Arabidopsis carotenoid cleavage dioxygenase 4 (AtCCD4) as regulator of carotenoid of seeds. | 7 | 6 | 77 | 6 | 13 | Ex, $x_0^\theta$ |
| Chen | The model by Chen *et al.* (2009) describes signaling in ErbB-activated MAPK and PI3k/Akt pathways, including seven receptor dimers and two ErbB ligands. | 500 | 3 | 105 | 4 | 191 | $E^{(1)}$, ev, NI, $x_0^{\text{fix}}$ |
| Crauste | The model by Crauste *et al.* (2017) describes CD8 T cell differentiation after virus infection. | 5 | 4 | 21 | 1 | 12 | Ex, NI, $x_0^{\text{fix}}$ |
| Fiedler | The model by Fiedler *et al.* (2016) describes Raf/MEK/ERK signaling in synchronized HeLa cells upon stimulation with MEK and ERK inhibitors. | 6 | 2 | 72 | 3 | 28 | $E^{(1)}$, NI, $u(t, \theta)$, $x_0(\theta)$ |
| Fujita | The model by Fujita *et al.* (2010) describes the epidermal growth factor (EGF)-dependent Akt pathway in PC12 cells. | 9 | 3 | 144 | 6 | 22 | Ex, ev, NI, $u(t, \theta)$, $x_0^\theta$ |
| Hass | The model by Hass *et al.* (2017) establishes early Reelin-induced signaling and identifies Src family kinases (SFKs) as crucial part for Dab1 signaling. | 9 | 6 | 221 | 23 | 62 | Ex, ev, $x_0(\theta)$ |
| Isensee | The model by Isensee *et al.* (2018) describes the Protein Kinase A (PKA)-II cycle in primary sensory neurons and its response to multiple stimuli, e.g., forskolin and cAMP analogues and is based on quantitative microscopy and Western blotting. | 25 | 3 | 713 | 109 | 46 | C, $E^{(1)}$, ev, NI, $u(t, \theta)$, $x_0^{\text{SSpre}}(\theta)$ |
| Lucarelli | The model by Lucarelli *et al.* (2018) describes activation of Smad proteins upon TGFβ stimulation, identifies the relevant complexes and linked them to target genes. | 33 | 43 | 1755 | 12 | 84 | $E^{(1)}$, Ex, NI, $x_0^\theta$ |
| Merkle | The model by Merkle *et al.* (2016) describes Epo-induced signaling simultaneously for CFU-E and H838 cells, with a parsimonious set of differing parameters. | 23 | 22 | 1141 | 62 | 197 | C, $E^{(1)}$, Ex, ev, $u(t)$, $x_0(\theta)$ |
| Raia | The model by Raia *et al.* (2011) describes interleukin-13 (IL13)-induced activation of the JAK/STAT signaling pathway for B-cells and two lymphoma cell lines. | 14 | 8 | 205 | 4 | 39 | C, $E^{(3)}$, $x_0^\theta$ |
| Schwen | The model by Schwen *et al.* (2015) describes binding of insulin to primary mouse hepatocytes based on flow cytometry and ELISA data. | 11 | 4 | 292 | 7 | 30 | $E^{(1)}$, NI, $x_0(\theta)$ |
| Sobotta | The model by Sobotta *et al.* (2017) presents IL-6-induced JAK1-STAT3 signal transduction and expression of target genes in hepatocytes. | 13 | 11 | 2220 | 110 | 269 | C, $E^{(1)}$, ev, $u(t, \theta)$, $x_0^{\text{SSpre}}(\theta)$ |
| Swameye | The model by Swameye *et al.* (2003), rapid shuttling of STAT5 from the nucleus back to the cytoplasm following Epo stimulus is recognized as a remote sensor. | 9 | 3 | 46 | 1 | 14 | C, Ex, NI, $u(t, \theta)$, $x_0(\theta)$ |
| Weber | The model by Weber *et al.* (2015) describes the interactions of PKD, PI4KIIIβ and CERT at the trans-Golgi network of mammalian cells. | 7 | 8 | 135 | 2 | 36 | $E^{(1)}$, ev, $u(t)$, $x_0^{\text{SSpre}}(\theta)$ |
| Zheng | The model is adapted from Zheng *et al.* (2012) and describes methylation at histone H3 lysines 27 and 36. | 15 | 15 | 60 | 1 | 46 | $E^{(1)}$, ev, NI, $u(t, \theta)$, $x_0^{\text{SSpre}}(\theta)$ |

The benchmark problems cover a wide range of model and data set sizes (Fig. 1A). A local identifiability analysis using the *identifiability test by radial penalization (ITRP)* (Kreutz, 2018) revealed that most benchmark models possess non-identifiable parameters. Furthermore, we found that initial conditions are specified in multiple ways, e.g., as equilibrium points of an unperturbed condition, and that different types of noise models and input functions are used (Fig. 1B). This results in a large number of combinations which have to be covered by computational modeling tools.

**Figure 1:**
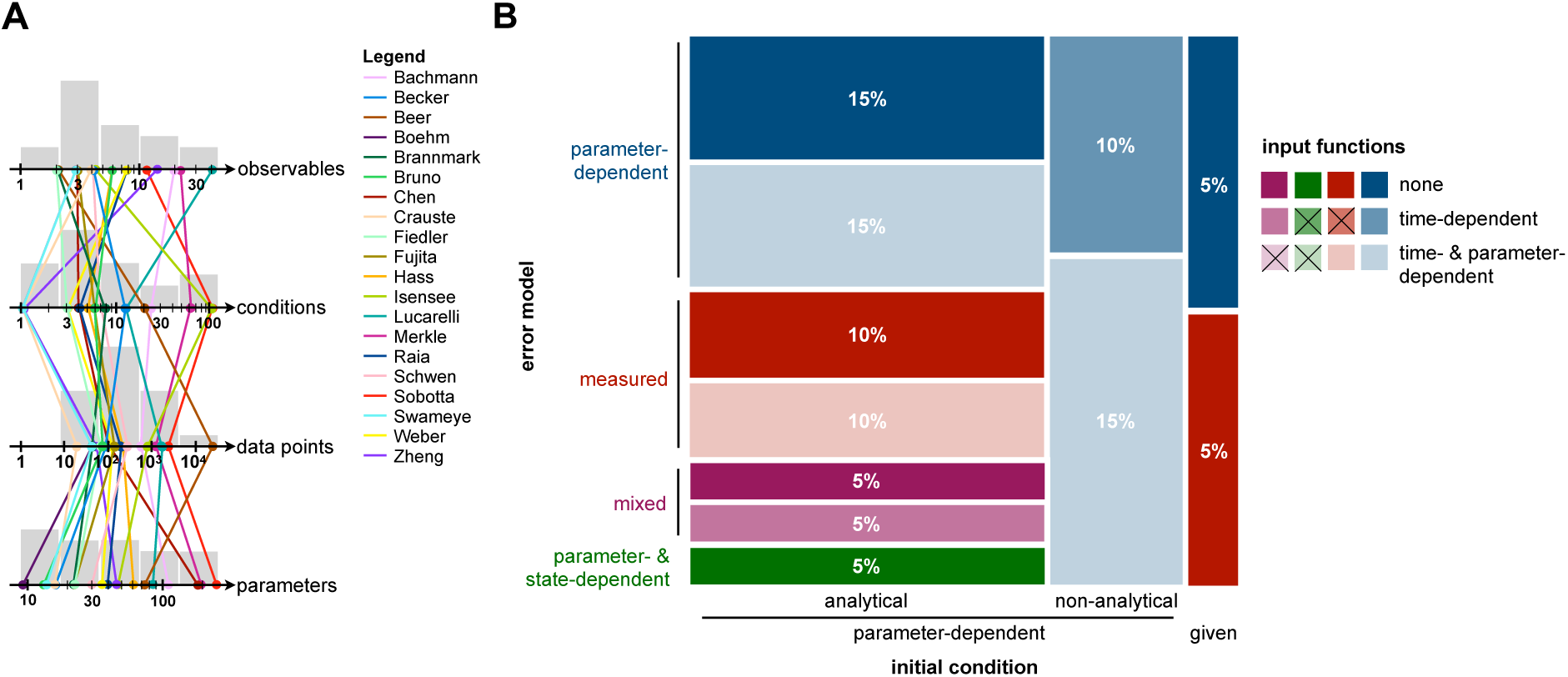
Property distribution in the presented benchmark collection. (A) Histograms for numerical model properties: number of observables, conditions, data points and parameters. Properties of individual models are indicated with an overlayed parallel coordinate plot. (B) Mosaic plot for the categoric model properties: initial conditions (columns), error models (color) and inputs (saturation). The areas encode the percentage of models with a particular combination of properties. Combinations of model properties which are not observed are crossed out in the legend. Nonanalytical parameter-dependent initial conditions can not be solved analytically and are obtained by simulating the system to steady state.

Although our collection is not unbiased, the spectrum of properties in the published models reveals requirements to be covered by modeling and parameter estimation tools. In the following, we will use the benchmark collection to assess a few common questions and statements.

### 4.2 Log-transformation of model parameters

A variety of studies in the systems biology field advocate the use of log-transformed parameters, ξ = log_10_(θ), for optimization:

> *“For parameters that are by definition non-negative a log-scale should be used in the parameter estimation.”* (Raue *et al*., 2013)

and recent evaluations verified that this can improve computational efficiency (Kreutz, 2016; Villaverde *et al*., 2018). A comprehensive evaluation on application problems is however missing and the precise reason for the improvement is still unclear. Here, we used the compiled benchmark collection to confirm the finding for multi-start local optimization (Fig. 2A) and to assess whether changes in the objective function landscape might be a potential reason. The performance metric is the average number of converged starts per minute (see Villaverde *et al*. (2018)). Starts are considered to be converged if the objective function value differs at most by 10^−1^ from the best objective function value found across all runs for the given benchmark problem, whereby we only included the models for which the best value was found more than once. We also found a strong dependence of the results on this threshold (see Supplementary Information, Fig. S9).

**Figure 2:**
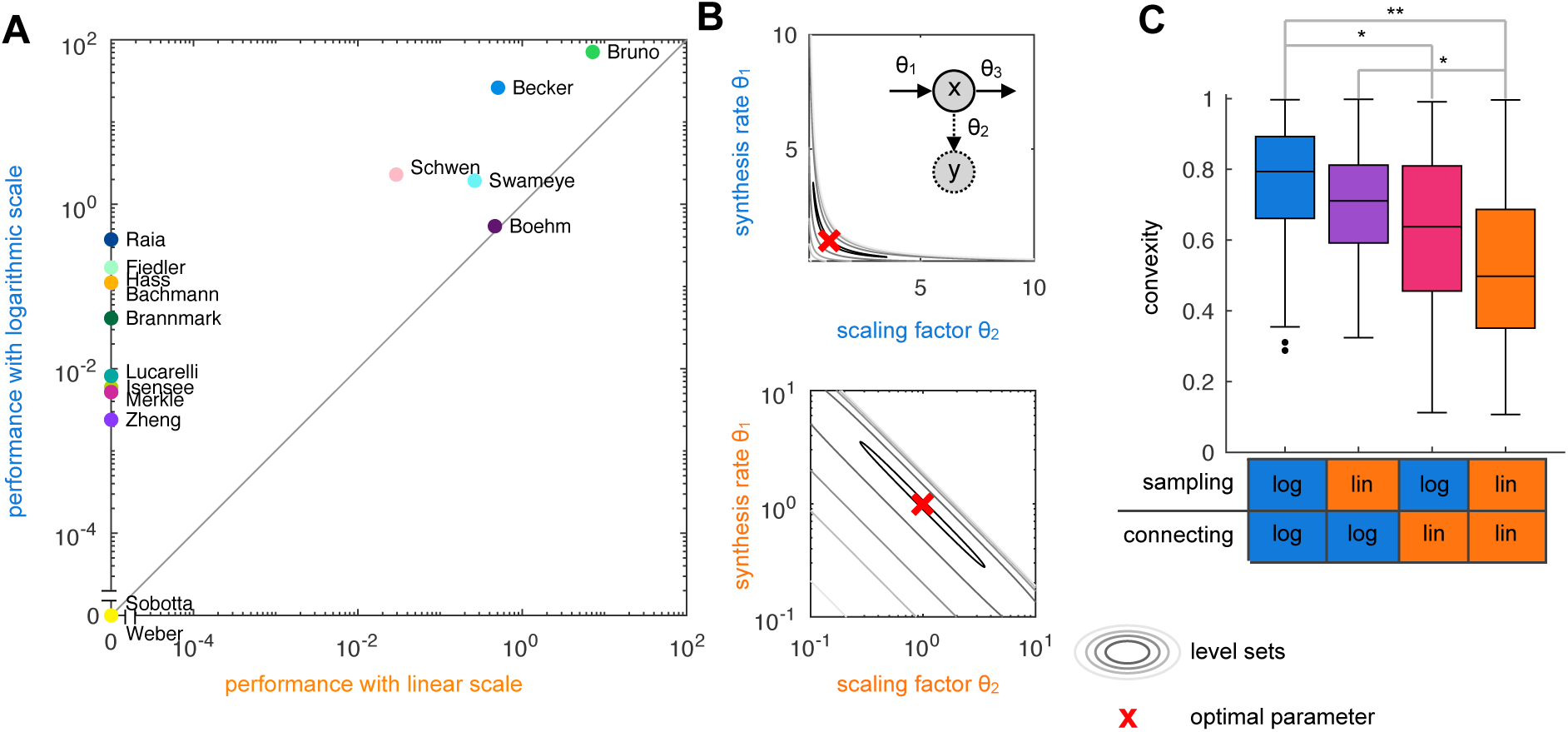
Linear vs. logarithmic scale. (A) Performance of the multi-start local optimization scheme using the MATLAB optimizer lsqnonlin for: (x-axis) sampling of initial values in log scale and optimization in linear scale; and (y-axis) sampling and optimization in log scale. Performance is measured as average number of converged starts per minute. (B) Level-sets of the objective function for a synthesis-degradation process (see Supplementary Information, Section 4) in linear parameters and log-transformed parameters. (C) Convexity properties of the benchmark problems in linear parameters and log-transformed parameters. It is indicated whether the two parameters are sampled in linear or log space and whether the connection between the two parameters is checked in linear or log space. Statistically significant differences are shown (p-value for rank sum test, * = < 0.05, ** = < 0.01).

Log-transformation leaves the optima unchanged but changes the shape of the level-sets of the objective function. We found several examples for which the level-sets are non-convex in the parameter *θ*, but convex in log-transformed parameters *ξ* (see, e.g., Fig. 2B). As local optimizers are well suited for convex problems, the change in the level set structure could be a reason for the improvement. To assess whether log-transformation improved the convexity of the objective function, we drew a random parameter vector *θ*^(1)^ *∈* Ω and a second random vector *θ*^(2)^ *∈* Ω with *‖θ*^(2)^ − *θ*^(1)^*‖* = 1 and a random location on the connecting line, *α ∼𝒰* (0, 1). For convex problems, the objective function *J* satisfies *∀ θ*^(1)^*, θ*^(2)^ and *α*:

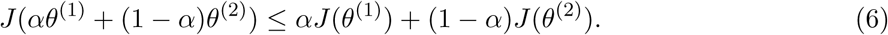

Accordingly, the fraction of triples (*θ*^(1)^*, θ*^(2)^*, α*) for which (6) holds provides a measure of convexity. We evaluated this measure for different combination of sampling strategies for *θ*^(1)^ and *θ*^(2)^ (lin or log scale, indicated in the x-axis of Fig. 2C), and checking the connecting the two parameters in lin or log scale (see Supplementary Information, Section 5). For each combination, we sampled 1000 triples. Our comparison revealed that for most application problems, log-transformation increases the considered measure of convexity (Fig. 2C). Indeed, some problems appear to be completely convex when using log-transformed parameters. This provides a mechanistic explanation for the observed improvement in optimizer convergence.

### 4.3 Performance of local optimization methods

The no free lunch theorem for optimization states that

*“[…] what an algorithm gains in performance on one class of problems is necessarily offset by its performance on the remaining problems.”* (Wolpert and Macready, 1997)

This implies that effective optimization relies on a fortuitous matching between an optimization method and an optimization problem. Here, we used the benchmark collection to assess the performance of the trust-region-reflective and the interior-point algorithm in the MATLAB function fmincon (The MathWorks, 2016) to parameter optimization problems encountered in systems biology. These local optimizers are widely used, indeed, there are studies using both optimizers to exploit there individual benefits and performance differences (Stapor *et al*., 2018). The choice of the optimizer has direct implication for multi-start local optimization methods (Raue *et al*., 2013) and meta-heuristics (Villaverde *et al*., 2018), but also for uncertainty analysis using profile likelihoods (Raue *et al*., 2009).

For fmincon, mainly the default settings provided by MATLAB were chosen, which can be obtained by optimoptions(^*’*^fmincon^*’*^). Therein, the algorithm was chosen as trust-region-reflective or interior-point, respectively. Additional changes to the default settings comprise:

- A user-defined gradient and Hessian for Gauss-Newton optimization.
- The tolerance on first-order optimality was set to 0.
- Termination tolerance on the parameters was set to 10^−6^.
- As subproblem-algorithm, *cg* was always chosen.
- The maximum number of iterations was set to 10000.

The trust-region-reflective algorithm is tailored to optimization problems with linear constraints. The trail step of the optimizer is obtained by minimizing a quadratic approximation of the objective function within the trust region (which is chosen adaptively). Parameter bounds are handled in the step construction by scaling and reflection. The interior-point algorithm is a general purpose method (and the MATLAB default) for optimization problems with linear and nonlinear constraints. It solves a sequence of approximate optimization problems with barrier functions. In each iteration a direct step obtained by solving the so-called Karush-Kuhn-Tucker condition or conjugate gradient step using a trust region is performed. For details we refer to the MATLAB documentation (The MathWorks, 2016).

We performed multi-start local optimization with 1000 fits for all benchmark models. Our results revealed that for the considered benchmark problems the trust-region-reflective algorithm tends to outperform the interior-point algorithm (Fig. 3 and Supplementary Information, Figs. S10S30). Indeed, the trust-region-reflective algorithm achieved a higher number of converged starts per computation time for 18 of the 20 benchmark problems and is for 9 benchmark problems the only algorithm finding the optimal solution. However, the optimal solutions for 2 benchmark problems were only obtained using the interior-point algorithm. Accordingly, although the trustregion-reflective algorithm (which is not the MATLAB default) achieves the higher reliability and performance, it can be beneficial to test alternative local optimizers. Additional information of the multi-start fits and the computation time for each model, as well as a comparison of the trustregion-reflective and the interior point method with the least-squares solver implemented in the MATLAB function lsqnonlin can be found in Supplementary Information, Sections 2, 6 and 7.

**Figure 3:**
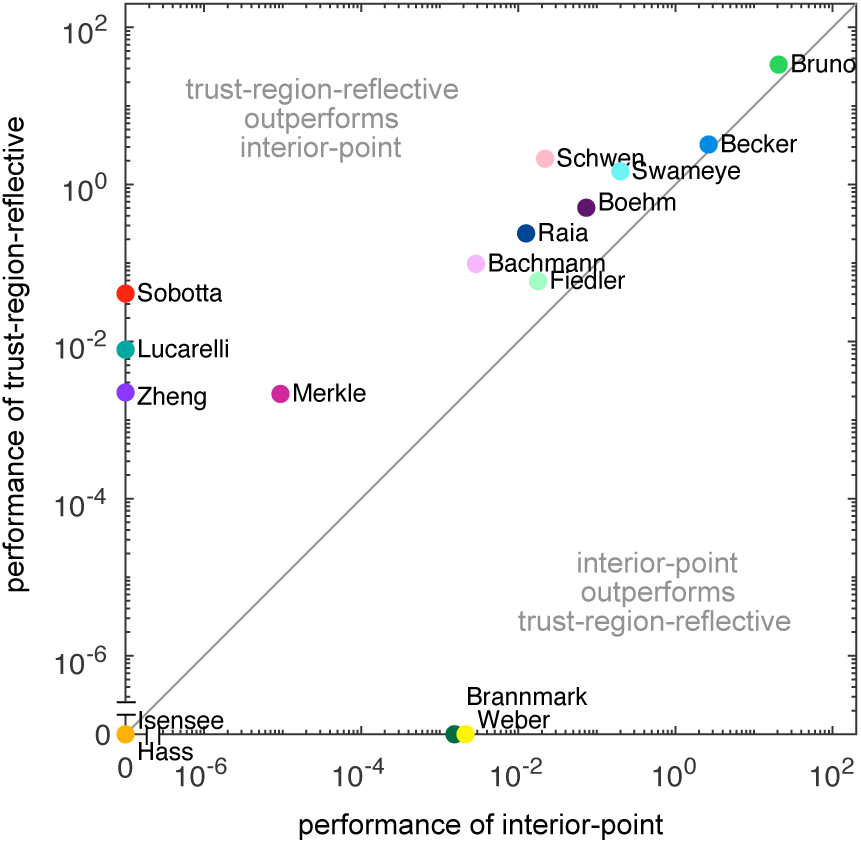
Comparison of optimizer performance. Scatter plot of the average number of converged starts per minute for the interior-point algorithm vs. trust-region-reflective algorithm.

### 4.4 Number of steps for local optimizers

Common questions in practical applications are (i) for how many steps (or iterations) a local optimizer should be run, and (ii) how the number of steps depends on the number of the parameters. For many local optimization algorithms, such bounds and results for scaling properties are available. For interior-point it has for instance been reported that for rather general classes of convex problems

> “[…] the number of Newton steps hardly grows at all with m [the number of constraints author’s note] (or any other parameter, in fact).” (Boyd and Vandenberghe, 2004, Section 11.5.6)

Similar findings are reported for other methods (see, e.g., Nesterov (2013)). As the independence of the number of optimization steps from the number of parameters might be surprising, we set out to assess the properties on the benchmark collection. For each problem, the trust-region-reflective algorithm implemented in the MATLAB function lsqnonlin was run, without constraints on the maximum number of function evaluation.

Our assessment of the average number of optimizer steps (Fig. 4) revealed that on average 391*±*19 iterations were taken. There is – as predicted by theory for convex problems – no significant dependence on the number of parameters (*ρ* = 0.02, p-value = 0.93). Accordingly, our analysis on the benchmark collection validated for the first time that the theoretical results also hold for application problems in systems biology (which are in general not convex).

**Figure 4:**
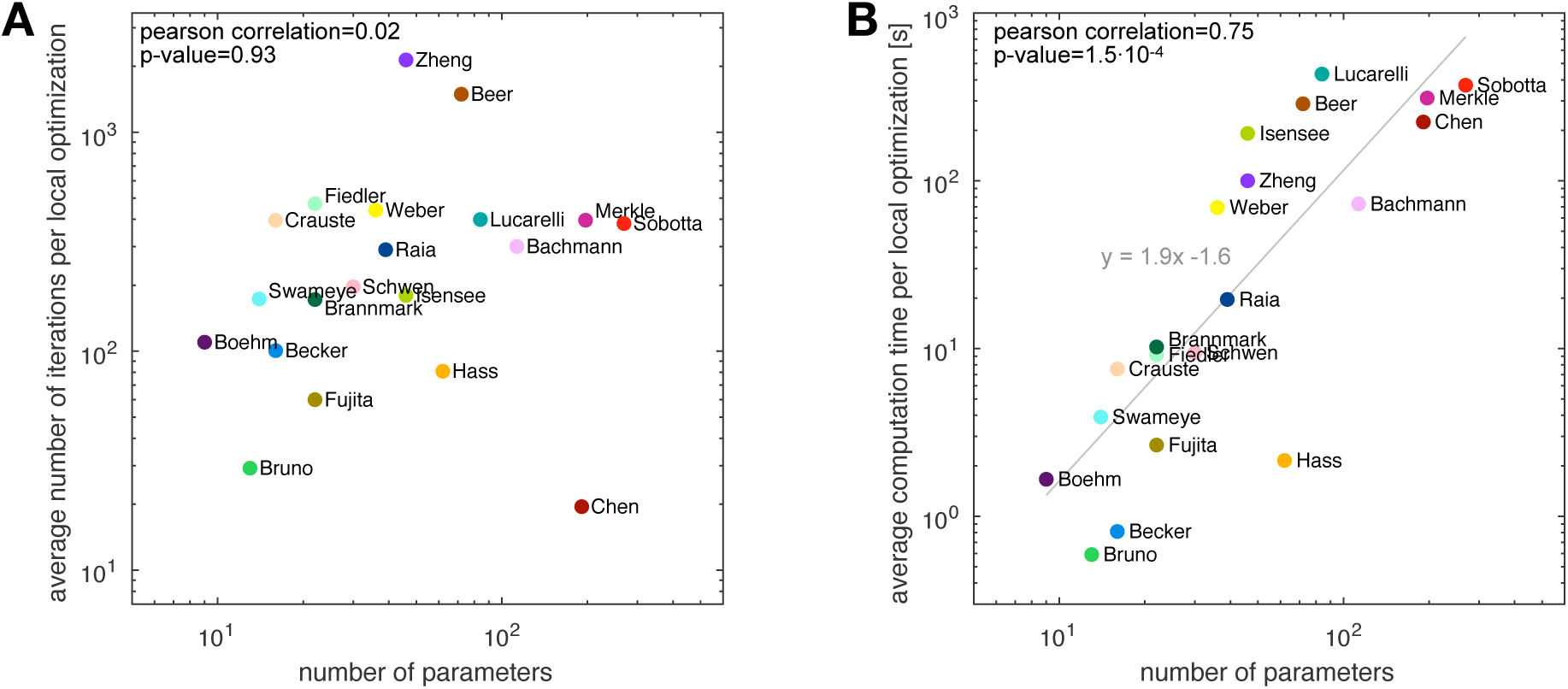
Influence of problem size. (A) Average number of optimizer iterations and (B) average computation time vs. the number of parameters. For optimization the trust-region-reflective algorithm implemented in the MATLAB function lsqnonlin was used and the averages across 1000 runs with different starting points were computed. The influence of the number of parameters was analyzed using correlation analysis and linear regression.

In contrast to the number of iterations, the computation time of local optimization depended on the number of parameters (*ρ* = 0.75, p-value = 1.5 *·* 10^−4^). For the trust-region-reflective algorithm using forward sensitivities for gradient calculation, we observed a roughly quadratic dependence 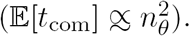

## 5 Discussion

Mechanistic dynamical models are used to describe and analyze biochemical reaction networks, to determine unknown parameters, gain biological insights and perform in-silico experiments. Novel methods to address these challenging tasks are proposed on a regular basis, however, a thorough assessment is often problematic. To address this problem, we compiled a collection of 20 benchmark problems. Reusability was ensured by providing the models in the machine-readable SBML format and the experimental data in structured Excel files. In addition, all aforementioned models are included in the open-source MATLAB toolbox Data2Dynamics (Raue *et al*., 2015) and the analysis scripts are provided as Supplementary Material.

To ensure that the benchmark problems are realistic and practically relevant, we exclusively included published models and measured experimental data. This is a key difference to existing benchmark collections which mostly considered models with simulated data (Villaverde *et al*., 2015; Ballnus *et al*., 2017). The benchmark models possess a broad spectrum of properties (e.g., different types of initial conditions, noise models and inputs), as well as challenges (e.g., structural and practical non-identifiabilities, and objective functions with multiple minima and valleys). The size of the benchmark problems ranges from roughly 20 data points, 10 parameters to be optimized and a single experimental condition to large models with more than 1000 data points, over 200 parameters and up to 110 distinct experimental conditions. This facilitates the assessment of the scaling behavior of novel algorithms.

We illustrated the value of the benchmark collection by performing three different analyses: (i) Our study of parameter transformations confirmed that optimization benefits from log-transformed parameter space. Furthermore, it suggested that the reason could be a significant increase of convexity of most problems, which provides a more benign setting for local optimizers. The observed change in the convexity appears to be the first mechanistic explanation for the observed improvement in optimizer performance. (ii) Our comparison of trust-region-reflective and interior-point algorithms revealed that the former is better suited for most parameter estimation problems encountered in systems biology. (iii) Our analysis of the scaling behavior confirmed theoretical results showing that the number of optimizer steps does not depend on the number of model parameters. The results of analyses (i)-(iii) could not have been obtained without the benchmark collection, which provided the means for a fair comparison. Indeed, the reliability of the findings depends directly on the size and the representativeness of the benchmark collection. Amongst others, previous studies were not able to provide an assessment of the scaling properties (Raue *et al*., 2013; Villaverde *et al*., 2015; Ballnus *et al*., 2017).

In conclusion, we think that the compiled benchmark collection will be an important resource for the systems biology community. It will facilitate the thorough evaluation of novel computational methods and support an unbiased assessment. In the future, the list of benchmark problems should be extended to enable a more fine-grained analysis and it should be integrated with public resources such as the BioModels database (Le Novere *et al*., 2006). Therefore, we encourage researchers to provide further models and data sets, e.g., by uploading them to our GitHub repository to obtain an even more powerful collection of benchmark models.

